# Microbial Community Structure of an Anaerobic Side-stream Coupled Anoxic-aerobic Membrane Bioreactor (AOMBR-ASSR) for an In-situ Sludge Reduction Process

**DOI:** 10.1101/2024.01.20.576432

**Authors:** Xinqiang Ning, Jialun Hu, Jiao Yue, Tang Tang, Bin Zhang

**Affiliations:** College of Bioengineering, Sichuan University of Science & Engineering, Yibin 644000, China; Chengdu Environmental Water Construction Co.Ltd., Chengdu Environment Group, Chengdu, 610000, China; School of Food and Biotechnology, Xihua University, Chengdu, 610039, China

**Keywords:** In-situ sludge reduction, anaerobic side-stream, high-throughput sequencing, microbial community

## Abstract

The in-situ sludge reduction process via the insertion of an anaerobic side-stream reactor into the sludge return circuit is an efficient approach to reduce the sludge yield in the activated sludge process. In this study, with the anoxic-aerobic membrane bioreactor (AO-MBR,CP) as a reference, high-throughput sequencing technology was used to reveal the characteristics of the microbial community structure in the anaerobic side-stream anoxic-aerobic membrane bioreactor sludge reduction process (AOMBR-ASSR,SRP). After the stable operation of two processes for 120 days, the average removal efficiencies of TN and TP in the effluent of SRP were increased by 5.63% and 29.85%, respectively. However, there was no significant difference between the two processes in the removal effect of COD and ammonia nitrogen. It is worth noting that the observed sludge yields (Y_obs_) of the two processes were 0.14 and 0.17 gMLSS/(gCOD), respectively, and the sludge reduction rate of the AOMBR-ASSR reduction process was 19.5%. Compared to the CP, the microbial richness and diversity index of SRP increased significantly. Among 14 major phyla, *Proteobacteria* and *Bacteroidetes* were the dominant microorganisms. *Chloroflexi*, which is responsible for the degradation of organic substances under an anaerobic condition, seemed to be reduced in the SRP. Meanwhile, other phyla that involved in the nitrogen cycle, such as *Nitrospirae* and *Planctomycetes*, were found to be more abundant in the SRP than in the CP. A total of 21 identified classes were observed, and primarily hydrolyzed fermented bacteria (*Sphingobacteriia, Betaproteobacteria, Actinobacteria*, and *Deltaproteobacteria*) and slow-growing microorganisms (*Bacilli*) were accumulated in the SRP. At the genus level, the inserted anaerobic side-stream reactor favored the hydrolyzed bacteria (*Saprospiraceae, Rhodobacter* and *Candidatus_Competibacter*), fermented bacteria (*Lactococcus* and *Trichococcus*), and slow-growing microorganisms (*Dechloromonas* and *Haliangium*), which play a crucial role in the sludge reduction. Furthermore, the enrichment of bacterial species related to nitrogen (*Nitrospir* and *Azospira*) provided the potential for nitrogen removal, while the anaerobic environment of the side-stream reactor promoted the enrichment of phosphorus-accumulating organisms.

## Importance

In the activated sludge process, the sludge recirculation line inserted into the anaerobic side stream reactor (ASSR) is considered to be a promising and cost-effective alternative process for in-situ sludge reduction. In the presence of ASSR, when sludge dissolves or hydrolyzes, the microbial community in the mainstream system will change due to changes in sludge characteristics. The difference in microbial community and substrate composition will lead to changes in the metabolic pathways of bacteria towards the substrate. Changes in metabolism may ultimately affect the growth of biomass, which is reflected in the growth of slow-growing microorganisms and the reduction of sludge production. reflected. Therefore, this study analyzes the microbial community structure in the side-flow reactor, which is helpful to reveal the relationship between microbial community structure and sludge reduction, and provides a theoretical basis for the application of anaerobic side-flow sludge reduction technology.

## 1. Introduction

Nowadays, the conventional activated sludge process is commonly used in the municipal sewage disposal, but the production of large amounts of excess sludge during the treatment process significantly restricts the sustainable development and application of this process (Khursheed, 2011; Cheng, 2017; Ferrentino, 2019). The high cost of residual sludge disposal accounts for 20%-60% of the total costs of wastewater treatment plants, and inadequate processing of the excess sludge treatment can easily cause secondary pollution to the environment (Niu, 2016a; Jiang, 2018). Therefore, considering the economic burden and the necessity of environmental protection, determining how to address the root of the problem of residual sludge has become a prevalent research topic for scholars both at home and abroad (Zhou, 2015a). Sludge reduction is divided into posterior sludge treatment/disposal and the in-situ reduction of sludge. In-situ sludge reduction is considered to be an ideal method to reduce excess sludge, which has less negative impact on process of sewage treatment (Meng, 2013).

Many studies have been carried out to develop of in-situ sludge reduction technology (Niu, 2016b; Habermacher, 2015). These technologies include dissolved oxygen (DO) and sludge retention time (SRT) controlls, adding chemicals to reduce biomass growth(Semblante, 2016). However, most of these methods led to an increase in operating costs and investment (Foladori, 2010). In addition, adding chemicals or using advanced oxidation processes may bring potential pollutants into sludge and sewage streams (Semblante, 2014). In-situ sludge reduction process with an anaerobic side-stream reactor (ASSR) added in the sludge return line has been considered as a promising way (Zheng, 2019c). Compared with other processes, ASSR process has such advantages as low operating cost, good process stability and easy management (Foladori, 2015). In the ASSR process, activated sludge (RAS) returned into the ASSR and then recycled into the main reactor of the process. Therefore, the sludge enters to alternating aerobic and anaerobic environmental cycle, and effective sludge reduction is achieved in the In-situ sludge reduction process (Jiang, 2018; Ferrentino, 2019). Many studies have been carried out to investigate the effects of oxidation reduction potential (ORP) (Saby, 2003), hydraulic retention time (HRT) (Jiang, 2018; Semblante, 2016), side-stream ratio of ASSR (Cheng, 2017) on sludge reduction, in order to minimize sludge production (Zheng, 2019c). According to the literatures, the sludge reduction efficiency (SRE) fluctuates greatly (0.2% - 60%) due to the difference of operation conditions and treatment water quality in the ASSR process. Therefore, it is important to explore the enhancement strategy to improve ASSR and ensure its high stability of SRE for its practical application (Zheng, 2019a; Pang, 2018).

Membrane bioreactor (MBR) technology, as a common wastewater treatment technology, can achieve high sludge concentration, sludge loading, and low sludge production (Liu, 2012). The combination of ASSR process and MBR technology is a promising in-situ sludge reduction method, which has the advantages of preventing the water degradagion, maintaining biological metabolism and having a small footprint (Cheng, 2017). The microbial population information of activated sludge is a vital functional index in the sewage treatment process (Li, 2014). In the sewage treatment process, the degradation of organic matter primarily depends on different microorganisms, and therefore the microbial population information is associated with the effect of sludge reduction and the removal of pollutants. Compared with traditional molecular biology methods (such as PCR-DGGE), high-throughput sequencing analysis can not only be utilized to comprehensively analyze the microbial community characteristics of the wastewater treatment process, but also to analyze the microbial flora associated with sludge reduction in the system (Hu, 2012).

In this work, using the anoxic-aerobic membrane bioreactor (AO-MBR) as a reference, the anaerobic side-stream anoxic-aerobic membrane bioreactor sludge reduction process (AOMBR-ASSR) was constructed. High-throughput sequencing analysis was used to analyze the microbial community structures of the two processes. The relationship between the microbial community structure composition and the sludge reduction was expected to be revealed. This study will provide a theoretical basis for the application of the AOMBR-ASSR.

## 2. Materials and methods

### 2.1 Experimental setup and operating conditions

The activated sludge used for this study was taken from the oxidation ditch of a sewage treatment plant (Zigong, China). In this study, two lab-scale membrane bioreactors (MBRs), namely an AOMBR for the control process (CP) and an anaerobic side-stream reactor coupled with AOMBR for the sludge reduction process (SRP), were both fed with synthetic wastewater with a constant flow rate of 48.96 L/d. The two systems were both stably operated for 120 d (Fig.1). In the CP, the effective volumes of anoxic, oxic, and membrane zones were 4.0, 13.5, and 4.7 L, with hydraulic retention times (HRTs) of 1.96, 6.62, and 2.3 h, respectively. An air diffuser was placed at the bottom of the aerobic reactor to maintain the dissolved oxygen (DO) concentration at 2.0∼4.0mg/L. The same air diffuser was placed at the membrane reactor to reduce membrane pollution. A mechanical stirrer was placed in the anoxic tank to mix the activated sludge.

**Fig. 1.**
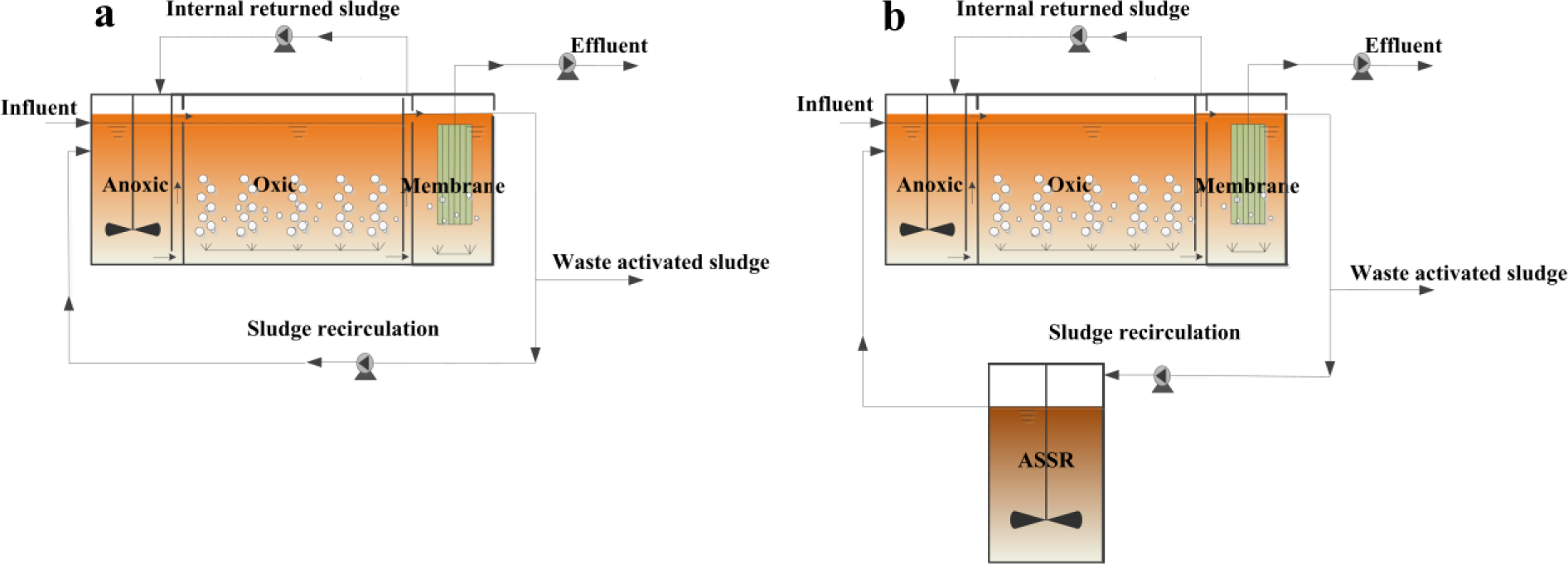
Schematic diagrams of the CP (a) and SRP (b)

The effective volumes of anoxic, oxic, and membrane zones in the SRP were the same size as the CP, but an additional ASSR was inserted into the sludge return circuit. The returned activated sludge (RAS) from the membrane reactor was pumped into an ASSR (14.6L volume) with a flow rate equal to 2.1 L/h, and then recycled to the anoxic reactor. Ratios of the internal returned sludge were both set to 300% for the two processes. The mixed liquor recirculation ratio of the CP was maintained at 100% of the influent flow rate. The side-stream recirculation rate of the SRP was controlled at 100% to achieve a recirculation ratio equal to that of the CP, which was consistent with previous studies (Coma, 2013).

### 2.2 Microbial Community Analysis

After the SRP and CP were stably operated for 90 days, sludge samples of the two processes were taken for microbial community structure analysis. According to the manufacturer’s agreement, genomic DNA of all samples was also extracted using the E.Z.N.A® Soil DNA Kit (Omega Bio-Tek, USA), and the content and quality of the extracted DNA were detected via 1% agarose gel electrophoresis. The V3-V4 region of the sample prokaryote 16S rRNA was selected for PCR amplification, and the primer sequences were 338F (ACTCCTACGGGAGGCAGCAG) and 806R (GGACTACHVGGGTWTCTAAT) (Niu, 2016a). The PCR amplifications were performed in a 20 μL reactor containing 4 μL of 5 × FastPfu Buffer, 2 μL of 2.5 mM dNTPs, 0.8 μL of each primer (5 mM), 10 ng of Template DNA, and 0.4 μL of FastPfu Polymerase (TransGen AP221-02, Beijing, China). The conditions for PCR were as follows: initial denaturation at 95 °C for 2 min, then 25 cycles at 95 °C for 30 s, 55 °C for 30 s, and 72 °C for 45 s; and a final extension at 72 °C for 10 min. After PCR amplification, the PCR product was detected via 2% agarose gel electrophoresis, and then recovered by the DNA Purification Kit (Axygen Biosciences, USA) and eluted with Tris-HCl.

After DNA extraction and PCR amplification, the amplified products were sequenced using the Illumina-MiSeq sequencing platform. The sequencing work was completed by Majorbio Bio-Pharm Technology (Shanghai, China). To obtain an effective sequence database for each sample, all the raw sequences were trimmed and removed for random sequencing errors and low quality sequences according to a previous method (Yuan, 2015). In order to fairly compare the two samples at the same sequencing depth, 39175 readings (with an average length of 436 bp) were normalized from each high-throughput output data value for further analysis.

MOTHUR software clustering was used to generate the operational classification unit (OTU) with a cluster similarity of 97%. OTU species identification was compared with the SILVA prokaryotic ribosomal sequence library (Release128 http://www.arb-silva.de), and with the confidence threshold set at 80%. Based on the clustering results, the following parameters were obtained from the four samples by the MOTHUR software: Chao and Ace, Shannon and Simpson, and Good’s coverage (Ning, 2014; Cheng, 2017).

### 2.4 Analytical Methods

Chemical oxygen demand (COD), total nitrogen (TN), ammonia nitrogen (NH_4_-N), nitrate nitrogen (NO_3_-N), and total phosphorus (TP) in the influent and effluent were analyzed every two days according to standard methods (Chinese NEPA, 2012). The suspended solids concentrations and volatile suspended solids concentrations of the mixed liquid in the CP and SRP were measured twice a week. Values of DO, pH, and ORP were monitored daily. One-way ANOVA was used to compare the differences between the pollutants in the effluents of the two processes. The observed sludge yield (Y_obs_, gMLSS/gCOD) was calculated by following the reported method (Sang, 2016).

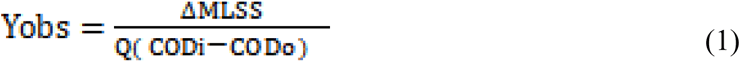

Where

*ΔMLSS* = the amount of excess sludge produced daily by the system (g/d),

*Q* = the treated water volume (L/d),

*CODi* = the system influent COD concentration (mg/L), and

*CODo* = the system influent COD concentration (mg/L).

## 3. Results and Discussion

### 3.1 Pollutant Removal Performances

The two processes were continuously operated for 120 days, and their pollutant removal performances are exhibited in Table 1. The average removal efficiencies of COD in the two processes were almost equally (95.3% VS 95.7%). The average removal rates of NH_4_-N in the CP and the SRP were 99.02% and 99.13%, respectively, and there was no significant difference between the two processes for the removal of NH_4_-N. Notably, the TN removal efficiencies of the CP and SRP were 76.39% and 82.02%, respectively, indicating that inserting the ASSR favored nitrogen removal (Pang, 2018). The average concentrations of TP in the effluent of CP and SRP were 1.76 mg/L and 0.65 mg/L, with corresponding removal efficiencies of 57.73% and 87.58%. Compared with the CP, the TP removal efficiency in the SRP was obviously improved, and will be associated with microbial community structure analysis in the later discussion. According to calculations, the Y_obs_ values of the CP and SRP were 0.17 and 0.14 gMLSS/(gCOD), respectively. Thus, the SRP achieved 19.3% of the sludge reduction effect compared to the CP. These results indicate that an ASSR promoted sludge stabilization and minimization .

**Table 1.** Average Characteristics of the influent and effluent wastewater of the CP and SRP process (mg·L^-1^)

| Items | COD | $\text{NH}_4\text{-N}$ | TN | TP |
| --- | --- | --- | --- | --- |
| Influent | $425.6 \pm 112.4$ | $52.64 \pm 11.64$ | $65.72 \pm 12.60$ | $4.22 \pm 0.77$ |
| CP effluent | $22.4 \pm 12.2$ | $0.48 \pm 0.73$ | $15.11 \pm 4.51$ | $1.80 \pm 0.89$ |
| SRP effluent | $19.6 \pm 12.0$ | $0.52 \pm 0.55$ | $11.10 \pm 3.31$ | $0.48 \pm 0.67$ |

### 3.2 Analysis of Bacterial Richness and Diversity

As exhibited in Table 2, the statistical results obtained from high-throughput sequencing illustrate that microbial community richness and diversity between the CP and SRP were different. In order to clearly understand the number of microbial communities, the sequence was clustering at a distance of 3%. The coverage index of sludge samples from the two processes was greater than 0.99 (*p* < 0.05), which indicated that the sequencing results could be considered to be effective in characterizing microbial community structure information. As can be determined by comparing Ace and Chao indices, the microbial community richness in the SRP reactor was significantly higher than that of the CP, which indicated that the inserted SSR had an pronounced effect on the microbial community structure. Additionally, the Shannon index not only characterizes the diversity of microbial community structures, but can also reflect the distribution of species (Zhou, 2015a). Therefore, the higher Shannon index illustrated the more diverse and even microbial communities in the SRP.

**Table 2.** Richness and diversity indexes of microbial communities of the CP and SRP process (α = 0.03)

| Sample | Reads | Normalization | OTUs | Ace | Chao | Shannon | Simpson | Coverage |
| --- | --- | --- | --- | --- | --- | --- | --- | --- |
| CP | 39175 | 10435 | 638 | 786 | 795 | 2.84 | 0.2916 | 0.995 |
| SRP | 45839 | 10435 | 824 | 912 | 921 | 4.45 | 0.0585 | 0.996 |

### 3.3 Analysis of Bacterial Richness and Diversity

As shown in Fig. 2, the relative contents of the bacterial communities were studied at different levels to investigate the phylogenetic diversity of the bacterial populations in the SRP and CP. 14 identified phyla were found both in the SRP and the CP (relative content > 1%) (Fig. 2). The most abundant phyla in the two processes were *Proteobacteria* and *Bacteroidete*s, which accounted for 64.16% and 15.23% in the SRP and 47.82% and 23.22% in the CP, respectively. In contrast to the CP, the relative content of *Proteobacteria* in the SRP was obviously reduced at the phylum level, and the relative content of *Bacteroidetes* was obviously increased, which indicated that the ASSR resulted in the extinction of some *Proteobacteria* in the SRP. This is primarily because most of the *Proteobacteria* are heterotrophic microorganisms; it is difficult for them to adapt to the ASSR, and some of them died out in the SRP. Due to the cell lysis of the *Proteobacteria*, the *Bacteroidetes* used intracellular substances released from the *Proteobacteria* for hydrolyzed fermentation and proliferation, and therefore the relative content of the *Bacteroidetes* in the CP was increased (Kim, 2009; Han, 2015). *Chloroflexi* is a common filamentous bacterium that are usually found in anaerobic environment of wastewater treatment processes (Kumar, 2010; Miura, 2008). *Chloroflexi* can degrade carbohydrates and cellular materials, and therefore they may play an important role in the SRP. In contrast with the average relative content of *Chloroflexi* (7.15% in the SRP and 11.89% in the CP) in the process, *Chloroflexi* seemed to be reduced in the SRP. Meanwhile, the relative contents of *Actinobacteria* and *Firmicutes* (5.88% and 4.80%) were respectively 2.58 and 2.46 times than those of the CP, indicating that they were significantly enriched in the SRP. *Actinobacteria* and *Firmicutes* were regularly associated with complex organic degradation; they could strengthen the removal capacity of pollutants in the SRP. Moreover, as a slow-growing microorganism with catabolism stronger than anabolism, the relative content of *Acidobacteria* in the SRP (1.51%) was higher than that in the CP (1.01%). Therefore, the enrichment of slow-growing microorganisms in the SRP would be cause sludge reduction. *Nitrospirae* is an important microorganism to realize nitrification during the wastewater treatment. It can be seen that *Nitrospirae* is obviously enriched in the SRP with a relative content of 2.64%, which indicates that the ASSR may improve the nitrification capacity of the SRP (Koch, 2015). The relative contents of the fermentative bacteria *Verrucomicrobia* and *Parcubacteria* were 1.54% and 1.25%, respectively, in the SRP, which were higher than those in the CP (0.66% and 0.17%). These bacteria contributed to the decomposition of intracellular substances in the ASSR. *Planctomycetes* is an anammox bacterium, and plays an important role in the conversion of nitrogen in the wastewater treatment process (Fuerst, 2011; Chistoserdova, 2004). The anammox bacterium is a type of auxoautotrophic bacteria, and can adapt to the special environment in the ASSR. It is enriched in the SRP, and the relative content of *Planctomycetes* is higher than that in CP.

**Fig. 2.**
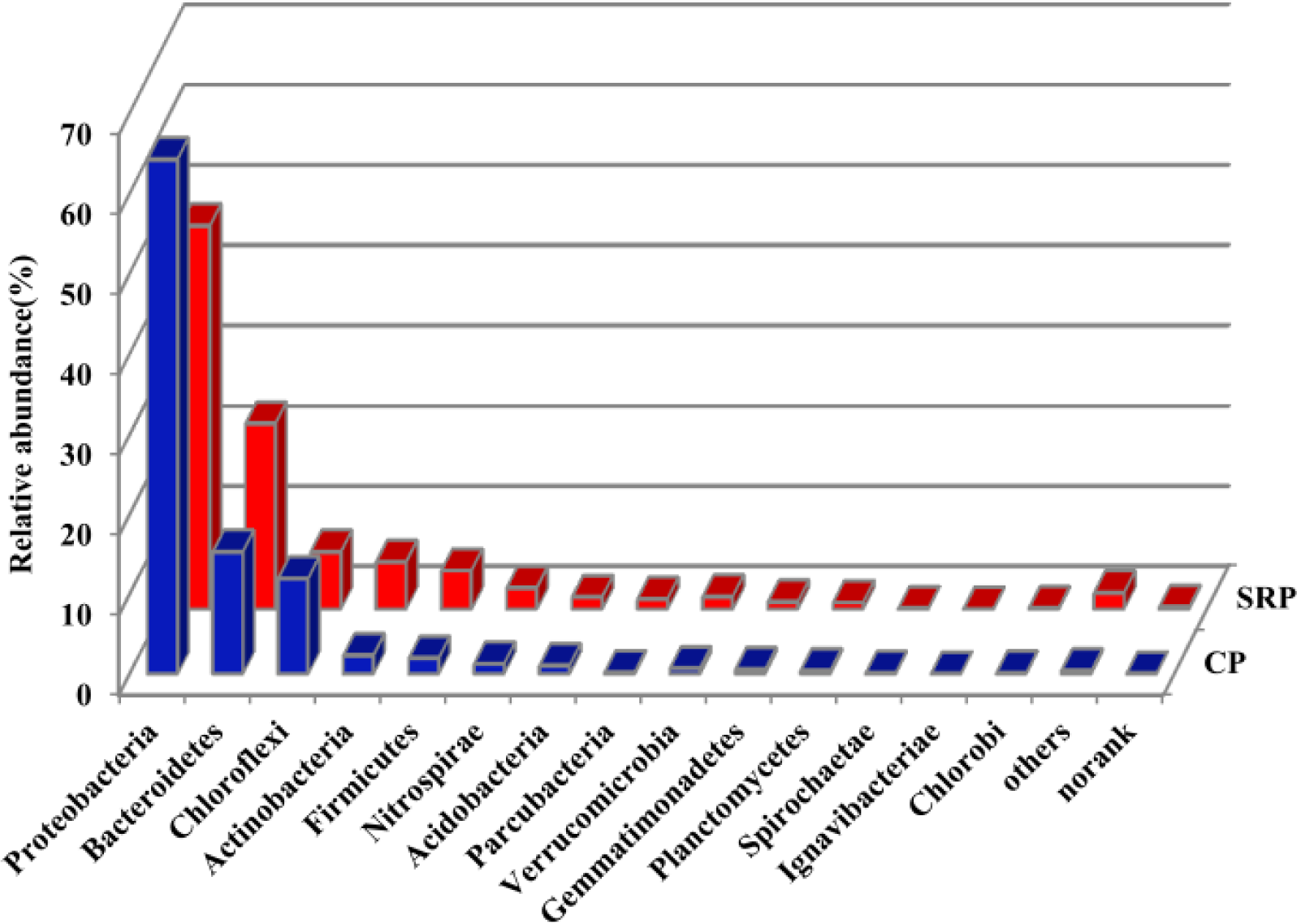
Phylum level

Further analyses of the microbial community information of the SRP and CP at the class level are provided in Fig. 3. A total of 21 identified classes were observed, and the *Gammaproteobacteria* was main subgroups, which belongs to the *Proteobacteria*. The relative content of *Gammaproteobacteria* was 54.66% in the CP and 25.12% in the SRP. This indicates that *Gammaproteobacteria* was reduced in the SRP due to the insertion of the ASSR. *Sphingobacteriia,* which can secrete extracellular hydrolase and conduct anaerobic hydrolyzed fermentation, was the second dominant bacterium in both processes (Zhou, 2015a). Its relative content was 19.99% in the SRP and 10.9% in the CP. Moreover, *Betaproteobacteria, Actinobacteria*, and *Deltaproteobacteria*, which are associated with hydrolyzed fermentation, were enriched in the SRP. The relative contents were 11.18%, 5.88%, and 4.58%, respectively, while those in the CP were 3.72%, 2.29%, and 2.68%. *Betaproteobacteria* mainly includes some sulfate reducing bacteria. A wide spectrum of carbon sources in the wastewater could be utilized by them (Niu, 2016b). The enrichment of such microorganisms in the SRP may be primarily related to the sludge reduction. *Alphaproteobacteria* and *Anaerolineae* are also anaerobic hydrolyzed fermentation bacteria that have a symbiotic relationship with methane-producing bacteria (Narihiro, 2012). The relative abundances of these bacteria in the SRP are respectively 2.3 and 2.7 times those in the CP. However, the relative content of *Cytophagia* in the SRP (1.62%) is lower than that in the CP (3.21%), which may be mainly due to its weak ability to adapt to the special environment of the ASSR as compared to other hydrolyzed fermented bacteria. *Bacilli* are not only a denitrification bacterium, but also a slow-growing microorganism (Jiang, 2018). Its relative content is 4.4% in the SRP and 1.7% in the CP. This indicates that its enrichment in the SRP would contribute to a higher denitrification capacity and a lower sludge yield.

**Fig. 3.**
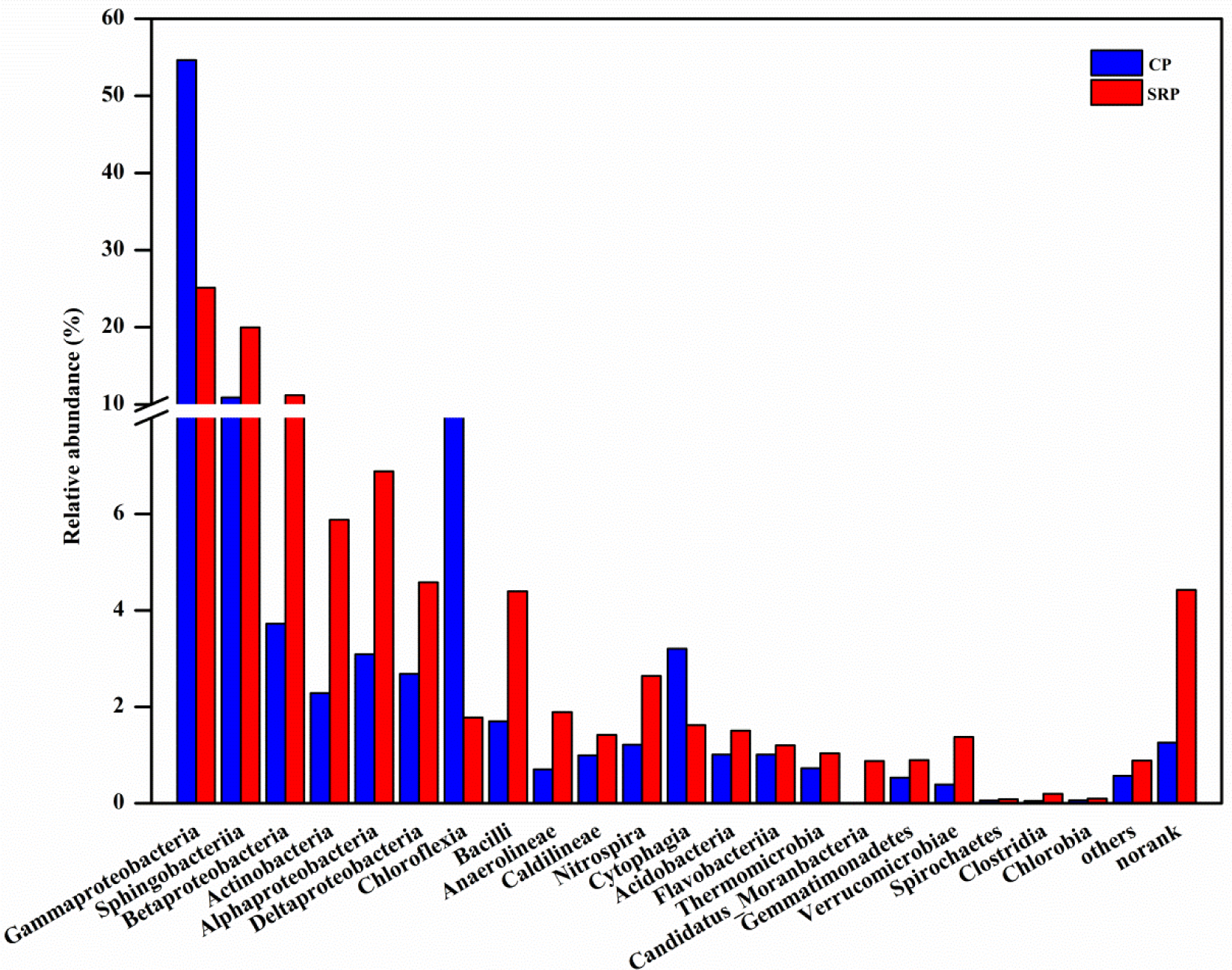
Class level

Bacteria of the SRP and CP on the genus level are shown in Fig.4. There are high relative contents of unclassified microorganisms on the genus level in the two processes. The unclassified microorganisms contributed to 14.83% (CP) and 25.59% (SRP) of the total genetic information. This indicates that the insertion of ASSR in the SRP prompted more unknown microorganisms that may be related to the specific conditions of the SRP. As shown in Fig.5, the differences of microbial community structure between CP and SRP on the genus level (the top 15 genera) are more intuitive, with the greatest differences being *Thiothrix*, *norank_f_Saprospiraceae*, *Roseiflexus* and *Dechloromonas*, respectively. *Thiothrix* and *Roseiflexus* are the most dominant strains. The relative content of CP was 53.05% and 8.96%, while that of SRP was 22.46% and 1.72%. These two microorganisms are commonly filamentous bacteria, and often lead to sludge bulking in the wastewater treatment process. Their reduction in the SRP may be related to the inhibition of sludge bulking due to ASSR insertion. *Norank_f_Saprospiraceae* can hydrolyze complex carbon sources and prey on other bacteria in the wastewater (Cheng, 2018), and the relative content of SRP (13.94%) is higher than that of CP (4.02%). *Dechloromonas* and *Haliangium* belong to the denitrifying bacteria (McIlroy, 2016; Ma, 2016). Their relative contents in the SRP (4.43% and 1.10%, respectively) were higher than those in the CP (0.85% and 1.09%). Because denitrifying bacteria usually grow slowly, *Dechloromonas* and *Haliangium* are slow-growing microorganisms, which were involved in the denitrification process (Liu, 2017). *Dechloromonas* is also a phosphate-accumulating organism (PAO) (Jiang, 2015). *Candidatus Accumulibacter*, a PAO, was also enriched in the SRP, and its relative content (0.34%) is about 1.54 times that in the CP (0.12%) (Xin, 2016). The relative contents of *Nitrospira* and *Ferruginibacter* bacteria, which are associated with nitrification in the wastewater treatment process, were 2.64% and 0.71% higher, respectively, than those in the CP (1.21% and 0.23%). Additionally, *Nitrospira* is the main nitrite oxidation bacteria, and can oxidize nitrite into nitrate by using hydrolysate in the SSR. This demonstrates that the ASSR is beneficial to nitrite oxidation bacteria growth, and favorable for the removal of nitrogen in the SRP. Moreover, the relative contents of *Lactococcus* and *Trichococcus* associated with anaerobic fermentation in the SRP were 3.06% and 1.24%, respectively, whereas in the CP they were 1.42% and 0.22%. Additionally, the enrichment of hydrolysis bacteria *Rhodobacter* and *Candidatus Competibacter* in the SRP (relative contents of 1.1% and 0.96%, respectively) revealed that the ASSR provided favorable conditions for sludge reduction with the action of hydrolyzed fermented bacteria.

**Fig. 4.**
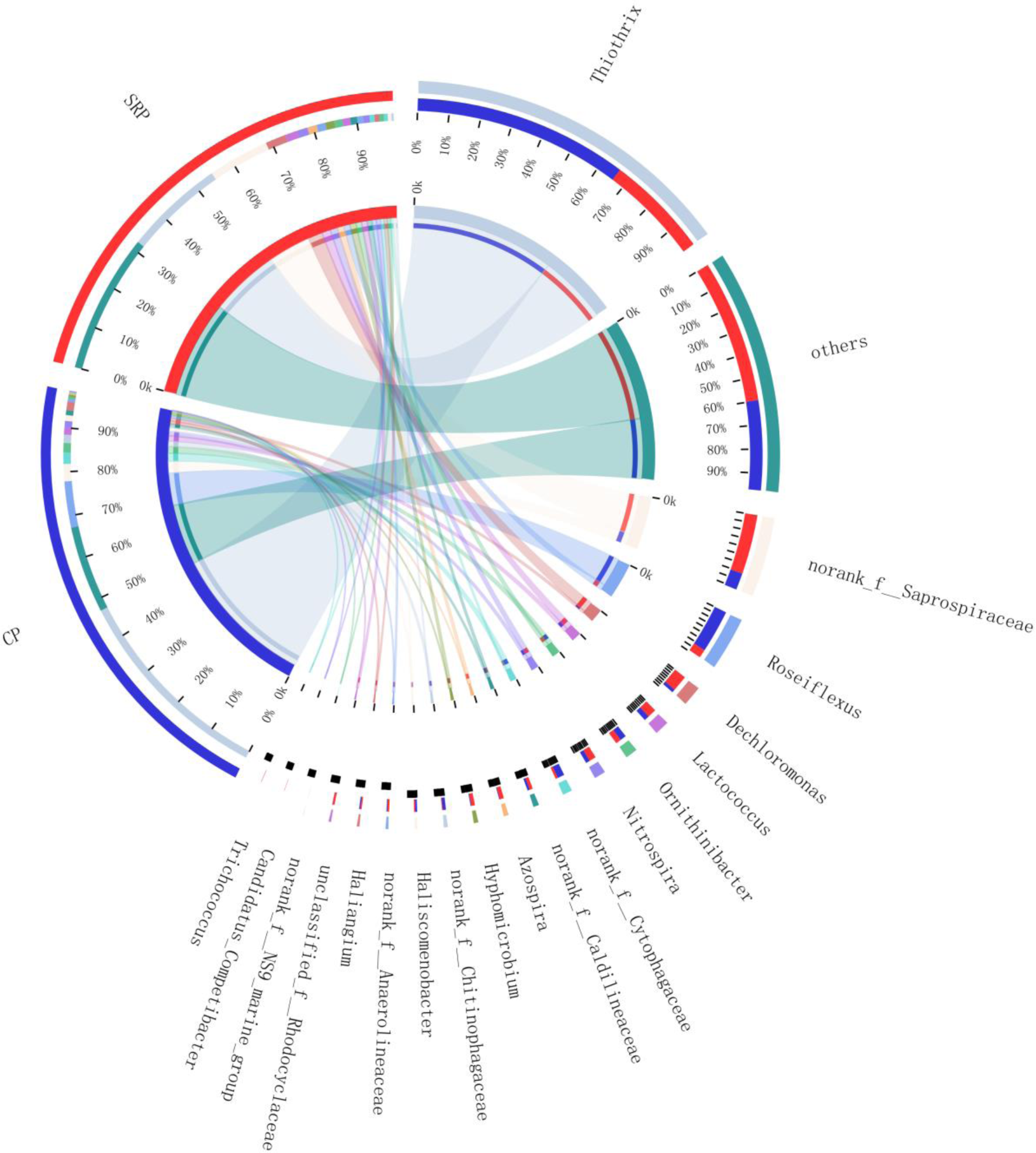
Genus level

**Fig. 5.**
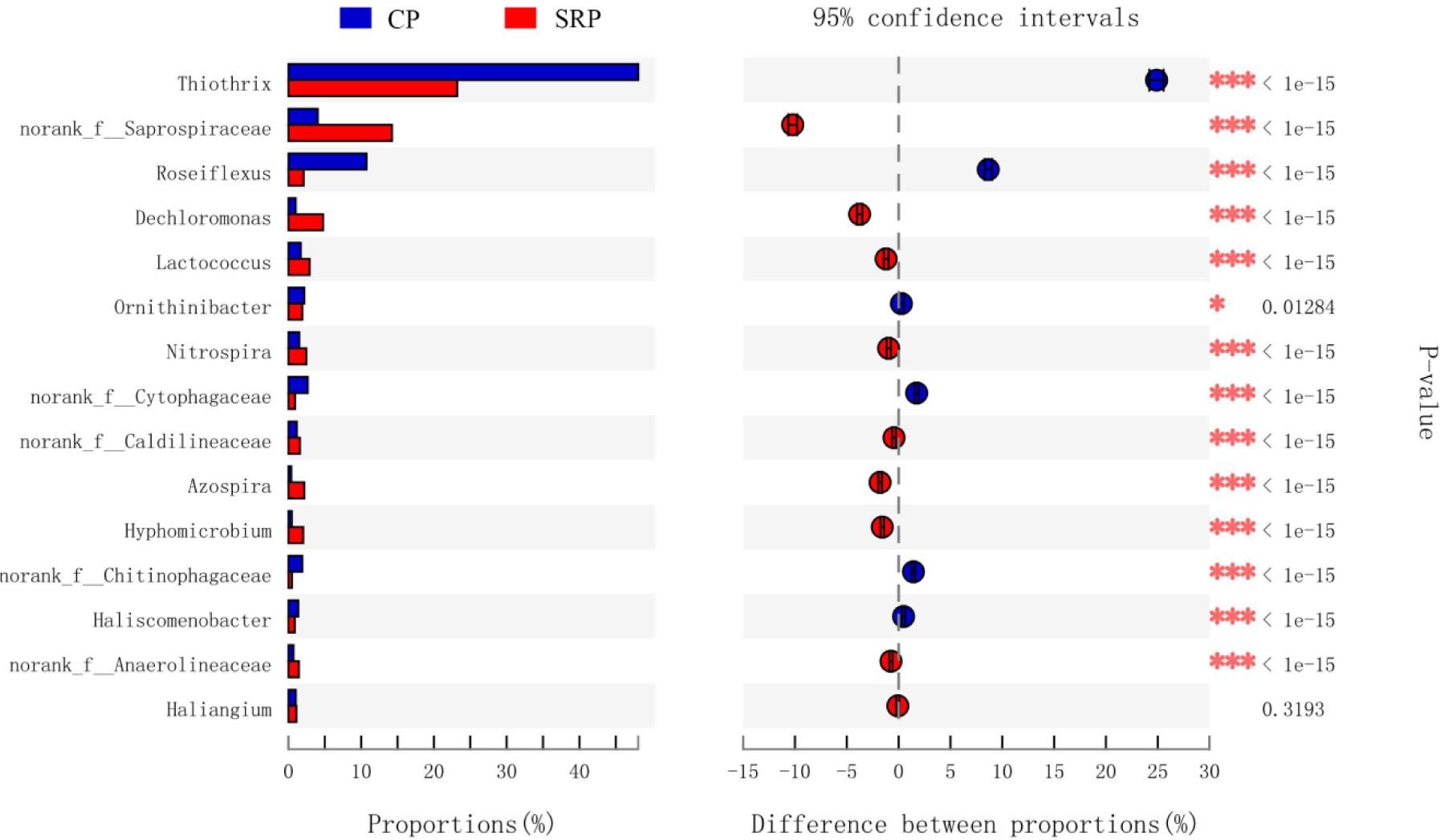
Bar plot of Wilcoxon rank-sum test at the genus level from the SRP and CP

### 3.4 Analysis of the relationship between microbial community structure and sludge reduction

The relevant existing literatures have confirmed that in-situ sludge reduction process could achieve excess sludge reduction without affecting performance of wastewater treatment (Semblante, 2016; Velho, 2016). A label-scale AOMBR-ASSR configuration completed by Saby (2003) sludge reduction efficiency varies from 23% to 58% with the Yobs 0.17-0.29 gTSS/gCOD. Zhou (2015b) implemented the same configuration in a pilot plant achieving 33% of sludge reduction (Yobs 0.21 gTSS/gCOD). Notably, in our study, the sludge reduction rate of laboratory-scale AOMBR-ASSR was 19.3% with the Yobs 0.14 gMLSS/gCOD, which lower than that of Saby(2003) and Zhou(2015b). The difference may be due to the difference of influent quality and sludge retention time of ASSR. In our study, sludge retention time in ASSR was 7.16h and high concentration synthetic wastewater was fed. However, in previous studies, the HRT(SRT) of ASSR was 10.4 h and real domestic wastewater.

It is suggested in the literatures that the mechanisms of sludge reduction that may be involved in ASSR include energy uncoupling, extracellular polymers (EPS) degradation, endogenous decay, and the select bacteria enrichment (Semblante, 2014; Oliveira, 2018). According to carbon balance analysis, sludge decay in ASSR has been proved to be the main reason for sludge reduction (Jia, 2013). However, sludge is composed of complicated bacterial biomass, and sludge decay is the result of various parts affected in ASSR system (Foladori, 2015). The anaerobic starvation environment in the ASSR affected the microbial community structure, and Foladori (2015) reported that the enrichment of hydrolyzed fermented microorganisms may be vital for sludge reduction. Zhou (2015) also found that the bacteria *Anaerolineae* and *Actinobacteria* play an important role in the A+OSA process, which are involved in hydrolysis and fermentation of different organic matter. Additionally, the enrichment of slow-growing *Trichococcus* could also favor sludge reduction. Cheng (2017) observed many hydrolyzed bacteria concentrations in the ASSR, while the nitrification bacteria *Nitrospirae* were mainly concentrated in the membrane reactor. In this study, the unique environment of the ASSR in the SRP promoted the growth of uncultured and unclassified microorganisms, which were capable of utilizing the substrate matrix returned to the main reaction zone. Diversity and richness of the microbial communities were likely improved. The enrichment of hydrolyzed and fermented bacteria and slow-growing microorganisms was responsible for sludge reduction. The hydrolysis process of granular organic matter is the limit step in the process of sludge reduction (Cheng, 2017), and therefore the enrichment of hydrolyzed bacteria, such as *Saprospiraceae*, *Rhodobacter*, and *Candidatus Competibacter*, at the genus level profited sludge reduction. Fermented bacteria such as *Lactococcus* and *Trichococcus* are essential in sludge reduction. In addition, slow-growing denitrification bacteria, such as *Dechloromona*s and *Haliangium*, reduced the sludge yield and enhanced the removal capacity of nitrogen. Furthermore, the enrichment of phosphorus-accumulating organisms such as *Dechloromonas* and *Candidatus_accumulibacter* in the SRP explained the obvious improvement of the phosphorus removal efficiency from a microbiological perspective.

## 4. Conclusions

In this study, the membrane bioreactor combined with the ASSR placed in the sludge return loop reduced the yield of sludge by 19.3%, and the average removal efficiencies of TN and TP increased by 5.63% and 29.85%, respectively. The results of high-throughput sequencing demonstrated that the diversity and richness of microorganisms in the reduction process were significantly improved. The enrichment of hydrolysis, fermentation, and slow-growth microorganisms was conducive to reducing sludge yield. The increase of the relative content of bacteria associated with nitrogen and phosphorus metabolism profited the removal of nitrogen and phosphorus. In future research, the analysis of microbial communities of the two processes will be critical to explore the possible functional microorganisms in the SRP and further improve sludge reduction efficiency of the ASSR.

## Acknowledgements

This research was supported by the Young Scientists Fund of the National Natural Science Foundation of China (51608339), the key SCI-tech project of Science and Technology Bureau of Zigong (2019YYJC09), College Students’ Innovative Entrepreneurial Training Plan Program (s201910622055), the Ministry of education ChunHui plan project (No. 191650[2018-93]), the Project of State Key Laboratory of Environmental Chemistry and Ecotoxicology, Research Center for Eco-Environmental Sciences, Chinese Academy of Sciences (No. KF2020-16), the Young Scholars Project of Xihua University in 2019 and the Department of Science and Technology of Sichuan Province (2017JY0129), China Postdoctoral Science Foundation (2019M650860), the Natural Science Foundation of Hebei Province (E2019402410), and Handan science and technology research and development program (No. 1623209044-2).

